# Active Transport of Membrane Components by Self-Organization of the Min Proteins

**DOI:** 10.1101/418384

**Authors:** YL Shih, LT Huang, YM Tu, BF Lee, YC Bau, CY Hong, HL Lee, YP Shih, MF Hsu, JS Chen, ZX Lu, L Chao

## Abstract

Heterogeneous distribution of components in the biological membrane is critical in the process of cell polarization. However, little is known about the mechanisms that can generate and maintain the heterogeneous distribution of the membrane components. Here we report that the propagating wave patterns of the bacterial Min proteins can impose corresponding steric pressure on the membrane to establish a directional accumulation of the membrane components, resulting in segregation of the components in the membrane. The diffusivity, influenced by the membrane anchor of the component, and the repulsed ability, influenced by the steric property of the soluble region of the component and molecular crowding, determine the differential spatial distribution of the component in the membrane. Thus, transportation of the membrane components by the Min proteins follows a simple physical principle, which resembles a linear peristaltic pumping process, to selectively segregate and maintain heterogeneous distribution of materials in the membrane.

## INTRODUCTION

Self-organization of biological molecules underlies fundamental cellular processes of all live forms. The Min system of *Escherichia coli* is a model system for studying protein self-organization that forms dynamic wave patterns on the membrane surface both *in vivo* and *in vitro*. The Min system, which consists of three proteins MinC, MinD, and MinE, mediates the cell division site placement [1]. The ATP-dependent interactions between MinD and MinE on the cytoplasmic membrane lead to cycles of pole-to-pole oscillation *in vivo*, thereby generating a concentration gradient of MinD that is high at both poles. Since the cell division inhibitor MinC interacts and oscillates along with MinD, the oscillation cycle facilitates division inhibition at the poles to prevent aberrant polar division [2-4]. Using the *in vitro* reconstitution approach, purified MinD and MinE in the presence of ATP can form propagating or standing waves on the supported lipid bilayers (SLBs) [5-7]. The formation of the MinDE wave patterns depends on the protein concentration, the molar ratio between MinD and MinE, and the geometrical confinement where the wave patterns are reconstituted [7-11]. Along with the experimental works, a series of numerical models have been reported to simulate the formation of the Min protein waves based on the reaction-diffusion theory [12-14].

Although the Min system have been well characterized and the protein-membrane interactions underlying the system is appreciated, whether the Min system can affect other membrane processes is not known. This brings up a notion that MinD belongs to the deviant form of the Walker-type ATPase family of proteins that has been implicated in partitioning cellular components in bacteria [15]. However, the division inhibitor MinC has been the only known cargo of the MinDE oscillator until now. Under the same theme to discover unanticipated functions of the Min system, a recent work reported that the Min oscillation cycle affected protein association with the inner membrane that resulted in modulating the cellular metabolism [16]. To follow up, we asked whether the Min system could act as a molecular machinery to spatially segregate membrane components in the membrane. In this study, we combined the experimental and theoretical methods to demonstrate that the propagating Min protein waves on SLBs can drive corresponding wave formation of the membrane components, leading to the spatial segregation of the membrane components. We conclude that the spatial distribution of the membrane components is determined by a balance between the steric repulsion caused by the Min proteins and the diffusivity of the components. This study may imply that pole-to-pole oscillation of the Min proteins inside a bacterial cell could selectively segregate membrane components to the designated membrane locations where the function of the component is required.

## MATERIALS AND METHODS

### Overexpression and purification of MinD

The His_6_-MinD fusion protein was expressed from BL21(DE3)/pLysS/pSOT4. A 24 mL overnight culture was used to inoculate 2.4 L fresh LB medium supplemented with 0.4% glucose, 34 µg/mL chloramphenicol, and 30 µg/mL kanamycin. The culture was grown in a 37°C shaker incubator until OD_600_ nm reached 0.4 to 0.6. The culture was cooled down to 16°C followed by addition of 0.7 mM isopropyl β-D-1-thiogalactopyranoside (IPTG) and cultured at the same temperature for an additional 16 hours. Cells were harvested and resuspended in 90 mL pre-chilled buffer A [50 mM potassium phosphate buffer pH7.4, 500 mM NaCl, 10% (v/v) glycerol, 5 mM β-mercaptoethanol, 2% Triton X-100, 3 tablets of cOmplete™, EDTA-free Protease Inhibitor Cocktail (Roche)] containing10 mM imidazole, followed by sonication on ice using a 1/2-inch (13mm) probe controlled by the Ultrasonic Processor VC750 (Sonics & Materials, Inc., USA). The cell lysate was centrifuged at 50,000 *×g* for 70 min at 4°C. The supernatant was filtered through a 0.22 µm syringe filter (Millex-GP, 33 mm, PES, Merck, USA) before being applied to a C10/10 Column (GE Healthcare Life Sciences, USA) packed with 5 mL His60 Ni Superflow resin (Clontech Laboratories, Inc.) at 4°C. After automated Superloop sample application, the column was washed with 100 mL buffer A containing10 mM imidazole, followed by a step gradient of imidazole from 10 mM to 90mM in a total volume of 150 mL. The protein was then eluted with 30 mL 400 mM imidazole and concentrated to 3 mL for gel filtration using a HiLoad 16/600 Superdex 200 pg column (GE Healthcare Life Sciences, USA) with the buffer containing 20 mM HEPES pH7.5, 450 mM KCl, 10% (v/v) glycerol, 0.1 mM EDTA, and 2 mM dithiotheritol (DTT). Fractions collected after gel filtration were examined using 12% sodium dodecyl sulfate polyacrylamide gel electrophoresis (SDS-PAGE) [17]. The protein concentration was determined using the Bradford Protein Assay kit (Bio-Rad Laboratories, Inc., USA). The protein sample was concentrated using 10KDa MWCO Amicon® Ultra Filters (Merck) when necessary before storing at −80°C.

### Overexpression and purification of MinE

An overnight culture of BL21(DE3)/pLysS/pSOT13 [P*T7::minE-his6*] [18] was diluted 40-fold into 1 L fresh LB medium supplemented with 0.4% glucose, 34 µg/mL chloramphenicol, and 30 µg/mL kanamycin. The culture was grown in a 37°C shaker incubator until OD_600 nm_ reached 0.4 to 0.6. The culture was cooled down to 16°C followed by addition of 0.5 mM IPTG and cultured at the same temperature overnight. Cells were harvested and resuspended in 25 mL pre-chilled buffer B [50 mM Tris-Cl pH7.4, 300 mM NaCl, 10% (v/v) glycerol] containing 5 mM imidazole, followed by three passages through the high-pressure cell disruption system at 30,000 psi to break cells. The cell lysate was centrifuged at 90,000 *×g* for 1 hour at 4°C. The supernatant was filtered through a 0.22 µm filter cup before applying to a 1 mL His-Trap HP Column (GE Healthcare Life Sciences, USA) at 4°C. The column was washed with 50 mL buffer B containing 40 mM imidazole. Protein was eluted with a linear gradient of imidazole from 40 mM to 300 mM in a volume of 30–50 mL. The peak fractions were pooled together and dialyzed against buffer C [25 mM HEPES pH7.6, 300 mM KCl, 10% (v/v) glycerol, 0.1 mM EDTA, 2 mM DTT], followed by concentration using 3K MWCO Amicon® Ultra Filters (Merck). The sample was further purified using a Superdex75 column (16/60; GE Healthcare Life Sciences, USA). The protein concentration was determined using the Bradford Protein Assay kit (Bio-Rad Laboratories, Inc., USA). The purified protein was examined by separation on 12% NuPAGE™; Bis-Tris Protein Gels (Thermo Fisher Scientific).

### Protein labeling

The amine-reactive dye, including the Alexa Fluor™ 594 NHS Ester (Thermo Fisher Scientific) and Alexa Fluor™ 647 NHS Ester (Thermo Fisher Scientific), was dissolved in dimethylsulfoxide (DMSO) to a final concentration of 10 μg/μL. Each 100 μL reaction contained 100 μg protein dissolved in PBS pH7.5, 100 mM sodium bicarbonate (pH 9.0), and 50–100 μg reactive dye. The reaction mixture was incubated with constant mixing in the dark at room temperature for 1 hour. The labeling reaction of MinE was passed through a Zeba^™^ spin desalting column (7K MWCO, Thermo Fisher Scientific) to remove excess dye, followed by exchange into the reaction buffer (25 mM Tris pH7.5, 5 mM MgCl_2_, 150 mM KCl) containing 10% (v/v) glycerol, aliquoted, and stored at −80°C. The labeling reaction of MinD was subjected to Sephadex G-25 fine buffer exchange before further purification by separation on a Superdex 200 Increase 10/300 GL column in the MinD gel-filtration buffer containing 10% (v/v) glycerol. The degree of labeling (DOL) was calculated according to the manufacturer’s instruction and was controlled between 0.5 and 1.5.

### Preparation of unilamellar vesicles for the formation of supported lipid bilayers

Phospholipids 1,2-dioleoyl-*sn*-glycero-3-phosphocholine 18:1 (δ9-Cis) PC; DOPC) and 1,2-dioleoyl-*sn*-glycero-3-phospho-(1′-*rac*-glycerol) (18:1 (δ9-Cis) PG; DOPG) were purchased from Avanti Polar Lipids (Alabaster, AL). N-((6-(biotinoyl)amino)hexanoyl)-1,2-dihexadecanoyl-sn-glycero-3-phosphoethanolamine (Biotin-X DHPE), Streptavidin Alexa fluor 488 conjugate (Alexa Fluor 488-Streptavidin), Lissamine™ Rhodamine B 1,2-Dihexadecanoyl-sn-Glycero-3-Phosphoethanolamine (Rhodamine DHPE), and 2-(4,4-Difluoro-5,7-Dimethyl-4-Bora-3a,4a-Diaza-s-Indacene-3-Dodecanoyl)-1-Hexadecanoyl-sn-Glycero-3-Phosphocholine (β-BODIPY™ FL C12-HPC) were purchased from Thermo Fisher Scientific Inc. (MA, USA). The lipid mixture was prepared with different composition, resulting in 67/33/0.5 mole% of DOPC/DOPG/Biotin-X DHPE, 67/33/0.25 mole% of DOPC/DOPG/Rhodamine DHPE, or 67/33/1 mole% of DOPC/DOPG/Bodipy FL C12-HPC. The mixture in chloroform was dried under nitrogen stream to leave a thin layer of the lipid cakes at the bottom of the container. The container was placed under vacuum to completely remove the solvent. The dried lipid cakes were rehydrated in the reaction buffer to a final concentration of 1 mg/mL. The rehydrated liposome solution was extruded sequentially through 400-nm, 100-nm, and 50-nm polycarbonate filters using an Avanti Mini-Extruder (Alabaster, AL) to prepare unilamellar vesicles. The extrusion was performed 19 times for each filter size. The vesicle suspension was stored at 4°C for up to 2 days.

### Preparation of microchannels

The Sylgard® 184 silicone elastomer kit (Dow Corning Corporation, California, USA) was used to prepare the polydimethylsiloxane (PDMS) channels. Briefly, the base (Sylgard 184A) was mixed thoroughly with the catalyst 87-RC (Sylgard 184B) at a 10:1 weight ratio, pulled into a 15-cm plastic Petri dish with a wafer of microchannel patterns placed on the bottom and degassed under vacuum more than an hour. The elastomer mixture was cured at 70°C for at least 3 hours to form a solid slab. The pieces were cut out from the solidified slab according to the microchannel design. The dimension of the microchannel was 1 cm × 1 mm × 16 µm (length × width × height) with circular reservoirs at both ends to accommodate liquid injection and removal during experiments (FIG. 1A). Holes were punched using a 14G flat-end needle at the reservoir position on both ends followed by cleaning with 95% ethanol and blow dried under a nitrogen gas stream. Immediately before assembly of the microchannel device, both the glass coverslip and the PDMS piece were subjected to surface treatment using the Expanded Plasma Cleaner (PDC-001, Harrick Plasma, New York, USA). Briefly, the coverslip was cleaned with argon plasma at 1.5 Torr for 10 min, followed by treating both the coverslip and PDMS piece with oxygen plasma using air at 400 mTorr for 75 sec.

**FIG. 1.**
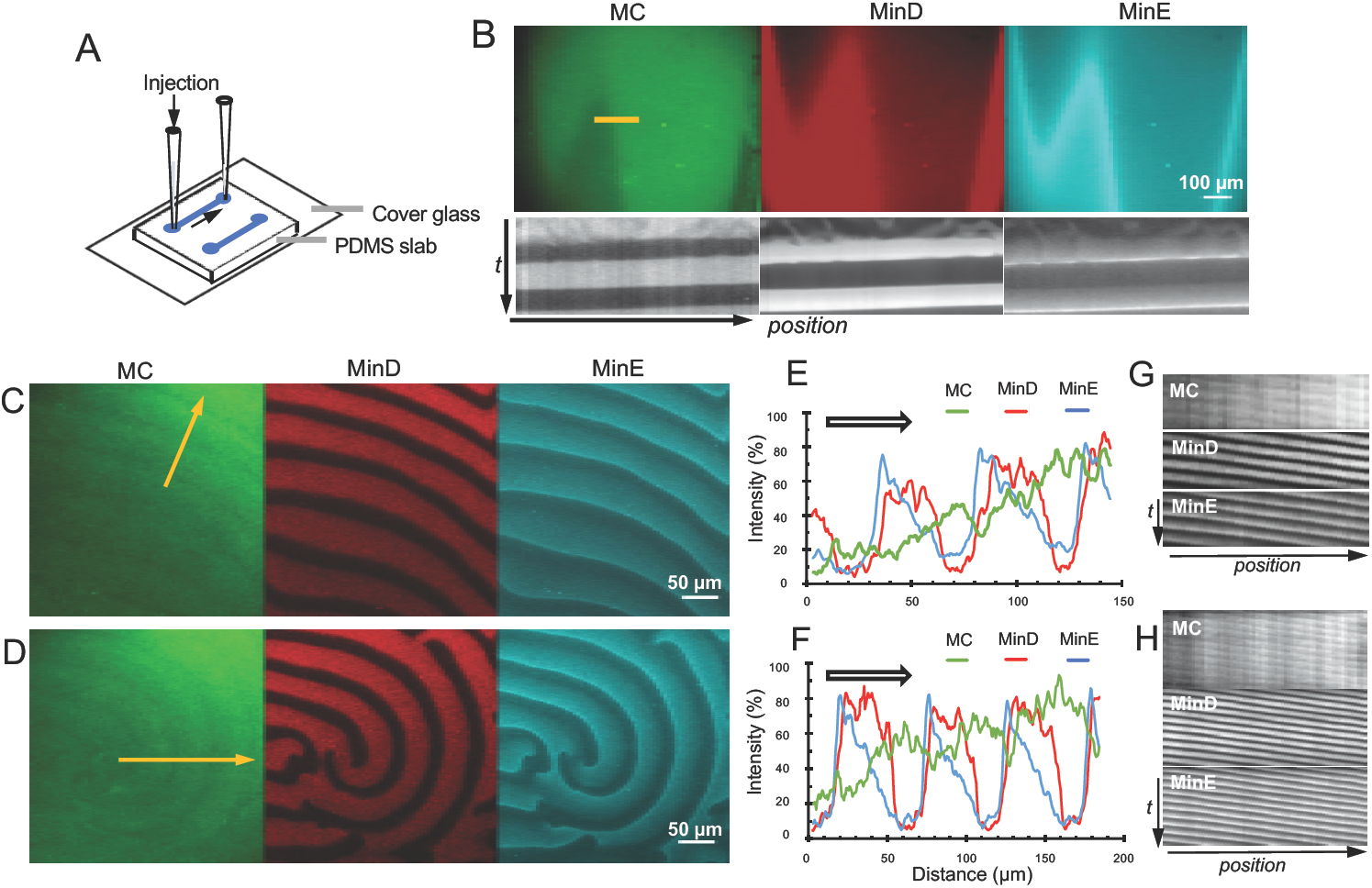
The corresponding membrane component (MC) waves of Streptavidin/Biotin-X DHPE formed along with the MinD and MinE protein waves. (A) Illustration of the microchannel for studying the membrane component waves. (B) The large-scale waterfall waves; Movie 1. (C) Traveling waves; Movie 2. (D) Spiral waves; Movie 3. (E, F) Intensity profiles and (G, H) kymographs of the Streptavidin waves. The intensity profiles and kymographs were obtained along the yellow arrows labeled in (B), (C), and (D). The arrows in C and D indicate the direction of wave propagation. The intensity profiles were measured from a single frame and partial kymographs were shown for clarity.

From the vesicle suspension, 50 µL was flowed into a microchannel for the formation of SLBs. The lipid suspension in the microchannel was incubated at room temperature for 1 hour. After incubation, the vesicle suspension was drawn out from the reservoir and replaced with 50 μL reaction buffer to wash off excess vesicles. For observation of the DOPC/DOPG/Biotin-X DHPE membranes, 1 µg/mL Alexa Fluor 488-Streptavidin prepared in the reaction buffer was filtered through a 0.1 µm syringe filter and incubated with the SLB at room temperature for 1 hour. The solution was removed followed by 5 washes with the reaction buffer. Before applying the reaction mixture, the microchannel was equilibrated with the reaction buffer containing 5 mM phosphoenolpyruvate (PEP) and 10 µg/mL pyruvate kinase (PK), and 5 mM ATP for 15 min.

### Reaction mixture of protein and membrane waves

The reaction mixture containing His_6_-MinD and MinE-His_6_ in a ratio between 1:1 and 1:1.2 was prepared in the reaction buffer without ATP, thoroughly mixed, and filtered through a 0.22 µm syringe filter. For observing the protein wave, 10–20 mole% of Alexa Fluor 594 His_6_-MinD and/or Alexa Fluor 647 MinE-His_6_ was included in the reaction mixture. We used the imaging conditions of rhodamine to observe Alexa Fluor 594 to avoid fluorescence leakage. The protein concentration varied slightly between experiments due to mixing in the fluorescently labeled MinD and/or MinE, but His_6_-MinD was around 10 µM and MinE was adjusted accordingly. 5 mM ATP was added to the filtered reaction mixture before applied to the SLB-coated microchannel for imaging waves under the microscope at room temperature.

### Fluorescence microscopy

Fluorescence images were acquired on the Olympus FLUOVIEW FV3000RS system equipped with the objectives UPLSAPO 10X2 (NA 0.4), UPLSAPO 20X (NA 0.75), and UPLFLN 40XO (NA 1.3), lasers of 488 nm (20 mW), 561 nm (20 mW), 640 nm (40 mW), and the high-sensitivity spectral detector (HSD). The spectral range to acquire images were set as 505–550 nm for Alexa Fluor 488, 570–620 or 630 nm for Alexa Fluor 594 and Rhodamine, and 650–750 nm for Alexa Fluor 647. The confocal pin hole was set large to allow enough light through for recording. The Olympus FV3/SW software was used for image acquisition. An inverted microscope (Olympus IX81, Olympus, Japan) equipped with a CCD camera (ORCA-R2, Hamamatsu, Japan), objective lens (UPLSAPO 20x, NA 0.75 and 10x, NA 0.4, Olympus, Japan), and filter sets for Alexa Fluor 488 (Chroma 49002), Cy3 (Chroma 49005), and Cy5 (Semrock Cy5-4040A) was also used. The Xcellence Pro software (Olympus) software was also used. Each microscopy image presented in the study represents a set of repeating experiments (greater than 3) using reaction mixtures prepared separately and were applied in different microchannels.

The acquired images were processed and analyzed using NIH ImageJ (Rasband, W.S., ImageJ, U.S. National Institutes of Health, Bethesda, Maryland, USA, http://rsb.info.nih.gov/ij/). The intensity profile was plotted using Microsoft Excel. The fitted trend lines are shown in the intensity profiles to facilitate visualization.

### Fluorescence recovery after photobleaching (FRAP)

The FRAP measurements were performed to measure the diffusion coefficients of membrane components. A 200 mW DPSS Blue Laser Modul (SEO, Taiwan) at 473 nm and Green Laser Module (Unice, Taiwan) at 532 nm were used to bleach a small region of interest (ROI) in SLB containing a fluorescently labeled membrane component for 0.2 sec. The intensity of a bleached ROI has a Gaussian profile with an approximate full width at half maximum (FWHM) of 10 µm. Recovery images were captured using the same wild-field fluorescent microscope setup as described above. The images were processed and the two-dimensional (2-D) diffusion coefficient of a membrane component in the membrane was calculated in MATLAB R2015a (Mathworks Natick, MA, USA) using an algorithm as previously described [19].

### Modeling the MinDE and MAC waves

A published model of the Min protein dynamics [14] sufficiently captured the characteristics of the *in vitro* Min protein waves, which further allowed us to correlate the membrane component dynamics with the existing model as reported in Results. The criteria that were used to build the kinetic model of the Min protein dynamics are as follows. (1) ATP is abundant in the system; (2) Because both MinD and MinE spontaneously form dimers in solution, the dimerization step is not included and MinD and MinE in the model refers to their dimeric forms; (3) Each MinD carries two MinE-interacting sites; (4) Either solution-form or membrane MinE can bind to one of the interacting sites on MinD to form a MinDE complex on the membrane; (5) Either solution-form or membrane MinE can bind to the empty interacting site on a MinDE complex which immediately causes dissociation of the entire complex with MinD going to the solution and MinE staying on the membrane; (6) MinD binds to the membrane with a rate constant *k*_*D*_; (7) *k*_*dE*_ is the rate constant for the solution-form MinE to interact with membrane-bound MinD or MinDE complex; (8) *k*_*de*_ is the rate constant for the membrane-bound MinE to interact with the membrane-bound MinD or MinDE complex; (9) *k*_*e*_ is the rate constant to described dissociation of membrane-bound MinE from the membrane. The concentrations, which change with time in the system are described as the following: *CD* (MinD in the solution), *C*_*E*_ (MinE in the solution), *C*_*d*_ (MinD on the membrane), *C*_*e*_ (MinE on the membrane), and *C*_*de*_ (MinDE complex on the membrane), respectively. The diffusivity of MinD, MinE, and the membrane component is described as *D*_*D*_ (MinD in the solution), *D*_*E*_ (MinE in the solution), *D*_*d*_ (MinD on the membrane), *D*_*e*_ (MinE on the membrane), *D*_*de*_ (MinDE complex on the membrane), respectively. Although the model considered several steps in bulk to reduce the complexity for simulation, it sufficiently reproduced the Min protein dynamic patterns. The kinetic model of the Min protein dynamics is shown in EQ.2–EQ.6. The concentration of membrane components (*C*_*m*_) that changes with time could be attributed by not only its own diffusion, but also the steric repulsion caused by MinD and MinDE complexes on the membranes, as shown in EQ.7.

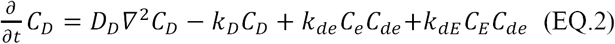

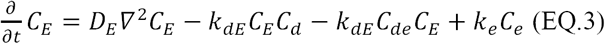

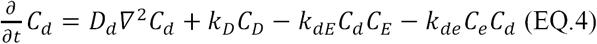

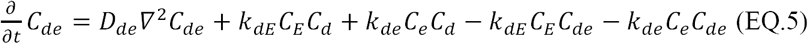

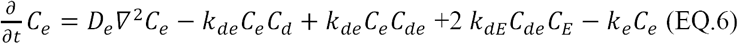

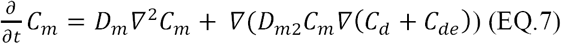

Comsol Multiphysics^®^ version 5.3a (Comsol Inc., MA, USA) was used to numerically solve the kinetic equations using 2-D spatial grids in a 300 µm × 150 µm region. The backward differentiation formula (BDF) method was used to solve the equations. Periodic boundary conditions were used for the top and bottom boundaries for all substances. While the periodic boundary conditions were also used from the two sides for the Min proteins in the solution, the no flux boundary conditions were set for the Min proteins attached to the membranes and for tested membrane components. The concentrations of MinD and MinE that were used in our experiments were 10 µM and 11.65 µM, respectively, and these concentrations are converted to 2-D concentrations by the method introduced by Loose et al. [5]. The diffusion constants of MinD and MinE in the bulk solution and the diffusion constants of the MinD, MinE, and MinDE complex on the membranes are based on the FCS and FRAP measurements reported previously [5]. The rate constants, including *k*_*D*_, *k*_*dE*_, *k*_*de*_, and *k*_*e*_, were derived from the literature [14]. The membrane component concentration is calculated from the molar ratio of the membrane component in SLBs.

## RESULTS

### Identification of the membrane component waves

To investigate whether the Min protein waves could influence the phospholipid bilayer, we adopted the *in vitro* reconstitution experiments [5] with emphases on the membrane components. We performed three-color, co-localization experiments of the membrane component, MinD, and MinE in microchannels (FIG. 1A) to obtain direct evidence for the existence of the membrane component waves. While MinD and MinE were conjugated with different fluorophores, the SLBs (DOPC:DOPG=67:33 mole%) were doped with 0.5 mole% Biotin-X DHPE that allowed subsequent treatment with Alexa Fluor 488 Streptavidin for examination of the membrane component waves. The reaction mixture of MinD, MinE and ATP was injected into the SLB-coated microchannels for studying the co-localization patterns of MinD, MinE and the membrane-tethered streptavidin.

Using this setup, we observed the streptavidin waves corresponding with the Min protein waves, indicating that the membrane component waves were generated alongside the Min protein waves (FIG. 1B-D, Movie 1-3). The protein and membrane component waves were dynamically evolving over time, showing up in different scales and wave forms. We described these initial large-scale waves as the ‘waterfall’ waves, because the spatial change of the fluorescence in all three channels resembled buckets of water pulling down the imaging field (FIG. 1B, Movie 1). The wavelength of the Min protein waves showed a wide distribution, but they were most frequently found between 50 and 60 μm with an approximate periodicity of 110 seconds (TABLE 1). Importantly, the observed Min protein waves showed characteristic features [5], i.e. the intensity profiles of both MinD and MinE waves were periodic, and the MinE intensity profile was asymmetrical with the intensity maximum spatially positioned behind the MinD waves at the trailing edge of the waves (FIG. 1E, F). Interestingly, the membrane component waves interdigitated with the Min protein waves and formed staggered spatial patterns on SLBs. The kymographs also demonstrated alternating spatial and temporal distribution of the Min protein and membrane component waves (FIG. 1G, H). Taken together, both staggered spatial and temporal patterns suggested that steric repulsion had occurred between the Min protein and membrane components during wave propagation. In addition, the membrane component waves were also recorded in the absence of the labeled proteins (FIG. 2A, B, Movie 4, 5); thus the observed membrane component waves were not caused by fluorescence leakage from other channels.

**TABLE 1.** Characteristics of the Min protein waves that were measured using the MinE intensity profile.

| Experiment | Label type | Wavelength<sup>c</sup> (μm) | Periodicity (sec) | Velocity (μm/sec) |
| --- | --- | --- | --- | --- |
| FIG. 1C | MinD AF568 | 54.07 ± 3.92<sup>d</sup><br>(n=5) | 109.33 ± 1.94<sup>d</sup><br>(n=9) | 0.522 |
|  | MinE AF647 |  |  |  |
|  | SLB AF488 S/B<sup>a</sup> |  |  |  |
| FIG. 1D | MinD AF568 | 51.79 ± 1.41<br>(n=6) | 115.50 ± 1.88<br>(n=8) | 0.448 |
|  | MinE AF647 |  |  |  |
|  | SLB AF488 S/B |  |  |  |
| FIG. 3A | MinD AF568 | 58.20 ± 2.03<br>(n=12) | 118.41 ± 1.98<br>(n=25) | 0.492 |
|  | MinE AF647 |  |  |  |
|  | SLB AF488 S/B |  |  |  |
| FIG. 4B | MinE AF647 | 119.48 ± 5.15<br>(n=8) | 189.16 ± 6.36<br>(n=19) | 0.632 |
|  | SLB Rhodamine |  |  |  |
|  | SLB Bodipy FL<sup>b</sup> |  |  |  |

<sup>a</sup> Alexa Fluor 488 Streptavidin/Biotin-X DHPE.
<sup>b</sup> Bodipy FL C12-HPC.
<sup>c</sup> The averaged wavelength corresponding to each Figure was measured from the same image series at different locations and/or different time frames.
<sup>d</sup> Standard error of the mean (SEM).

**FIG. 2.**
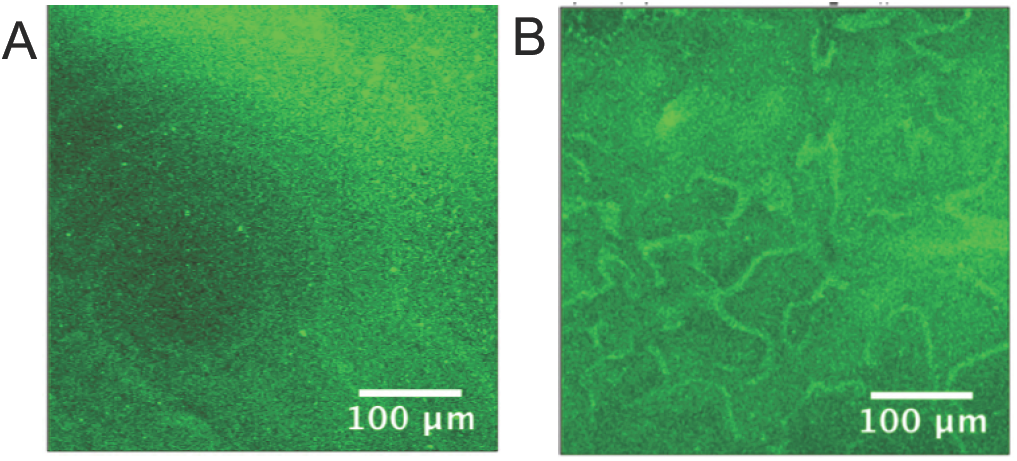
The membrane-tethered streptavidin waves were observed in the reactions without labeled Min proteins; Movie 4, 5.

### The membrane component accumulated in the same direction as the Min protein wave propagates

A striking feature associated with the membrane component waves was the long-range fluorescence intensity gradients. As shown in FIG. 3 and Movie 6, the fluorescence intensity of the Streptavidin/Biotin-X DHPE ascended from the center outwards in the spiral waves, which was specific to the membrane component and not found in the protein channels. The directional accumulation of the membrane component was also presented in the kymographs and the overall trend of the intensity profiles (FIG. 3B, C). Since the concentration of the fluorescent streptavidin used in the experiment was below its quenching limit, the fluorescence intensity was expected to be proportional to the concentration of the membrane component and the intensity distribution should represent the concentration distribution of the membrane component. Therefore, propagation of the Min protein waves could not only drive membrane components to migrate along with the proteins, but also led to the establishment of a concentration gradient of the membrane component in the membrane.

**FIG. 3.**
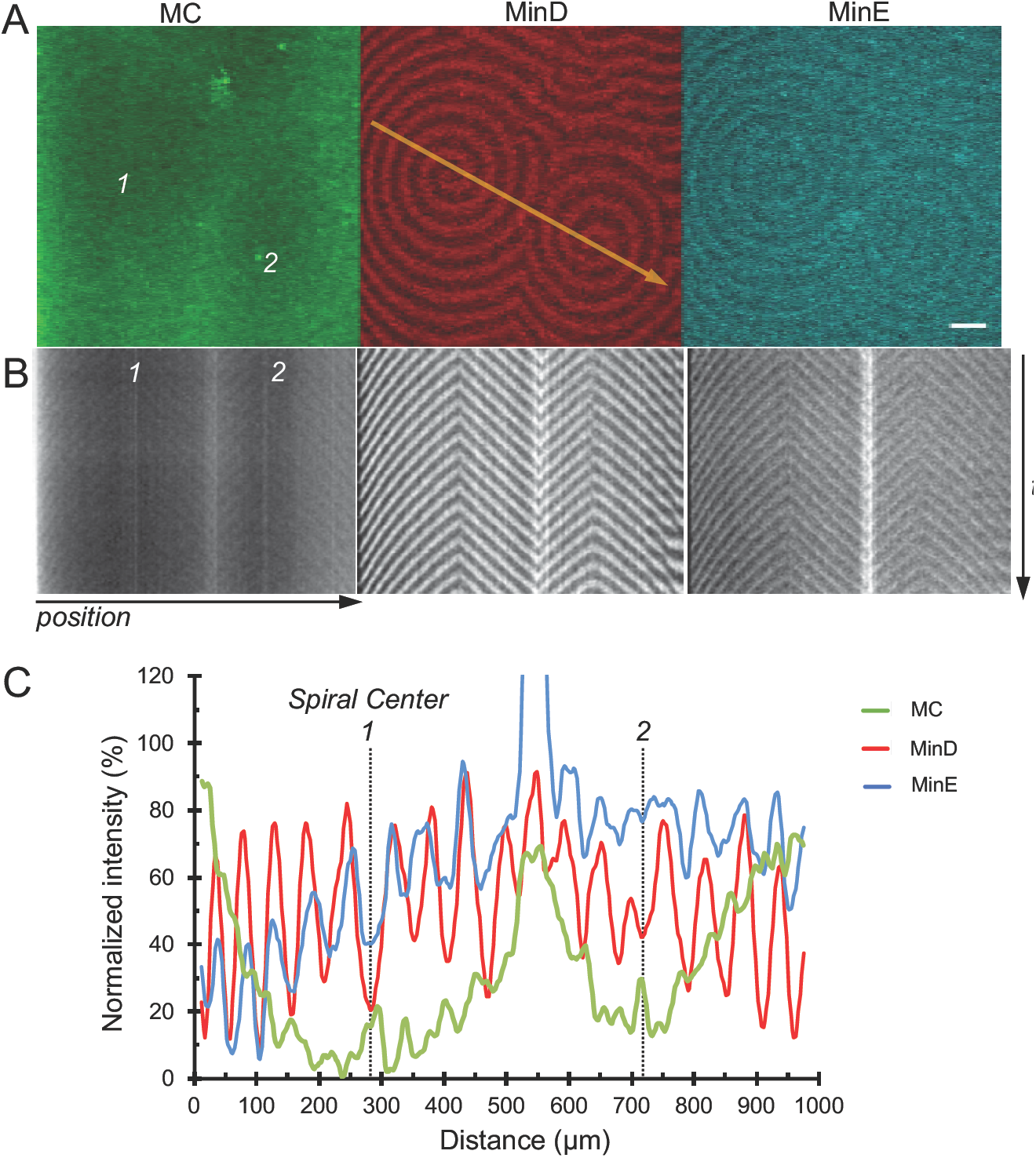
The concentration distribution of the Streptavidin/Biotin-X DHPE within spiral waves. (A) Confocal micrographs demonstrating the streptavidin, MinD, and MinE waves. The spiral centers are marked with 1 and 2. Scale bar, 100 µm. See Movie 6. (B) Kymographs and (C) spatial profiles were measured along the yellow arrow in (A).

### The steric property of membrane component contributed to the wave formation

In order to explain the steric repulsion between the membrane component and the Min proteins that led to the formation of the membrane component waves, we hypothesized that the soluble region of a membrane component could be the region that faced the repulsive force coming from the Min proteins to induce fluxes of the membrane component. The hypothesis predicted that a larger soluble region would face larger repulsive force, which could facilitate formation of the induced flux and establish a directional accumulation of the membrane component. The soluble region of a membrane component referred to the region above the phosphate group of the phospholipid headgroup and was exposed in solution (FIG. 4A).

**FIG. 4.**
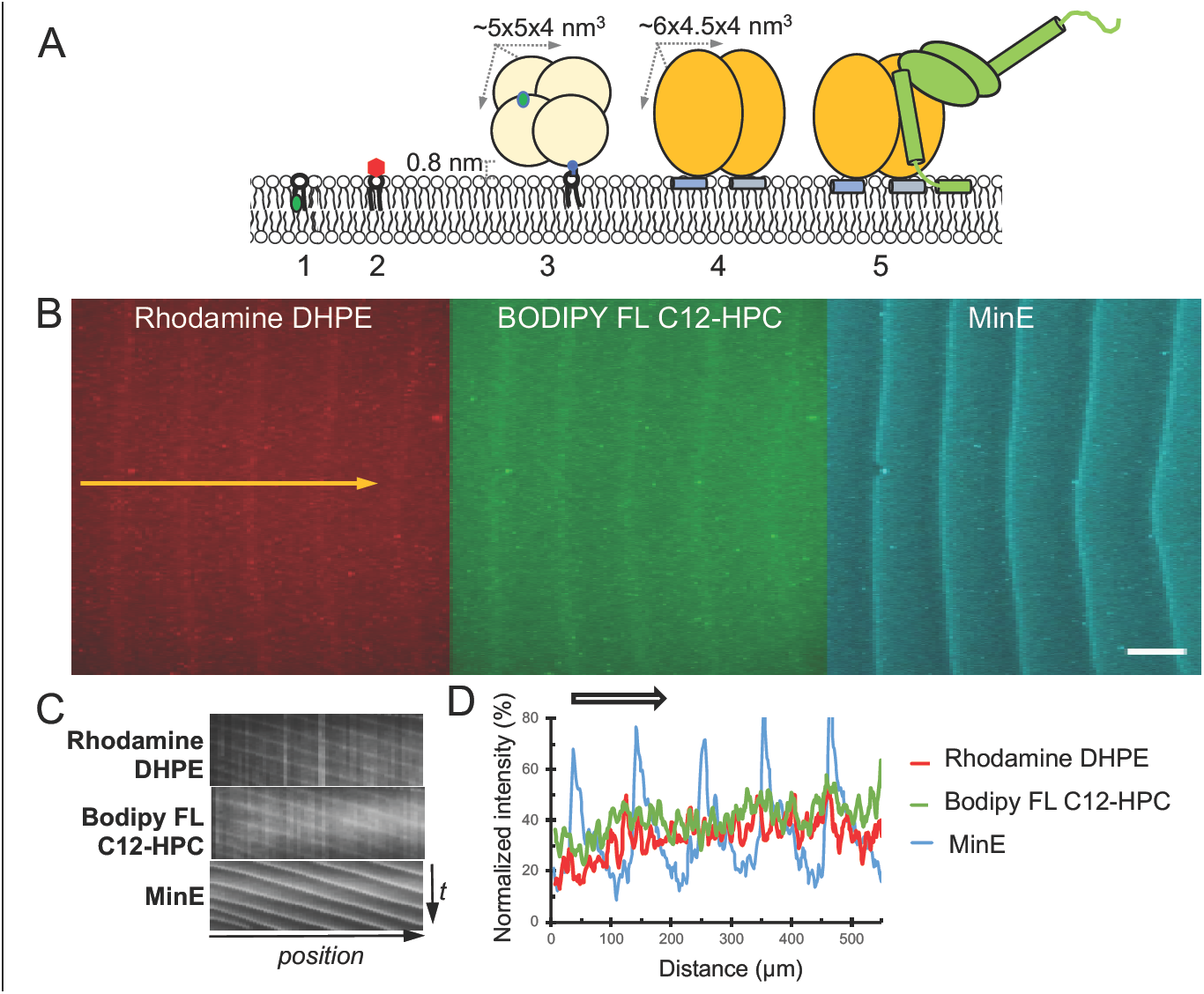
The steric property of the membrane component is critical for the wave formation. (A) Illustration of the steric difference between different membrane components, MinD dimer, and MinDE complex. 1. BODIPY FL C12-HPC; 2. Rhodamine DHPE; 3. Biotin-X DHPE/AF488 Streptavidin; 4. MinD dimer on membrane; 5. MinD-MinE complex on membrane. (B) The waves of both Rhodamine DHPE and Bodipy FL C12-HPC were observed simultaneously along with the MinE waves, but no spatial exclusion between them and no directional accumulation of the components was identified. Scale bar, 100 µm. See Movie 7. (C, D) Kymographs (C) and spatial profiles (D) of Rhodamine DHPE, Bodipy FL C12-HPC, and MinE. The kymographs and spatial profiles were obtained along the yellow arrow labeled in (B). The arrow in (D) indicates the direction of wave propagation.

To examine the hypothesis, we studied the headgroup-labeled Rhodamine (360.4 Da) and the fatty acid tail-labeled BODIPY FL C12-HPC, which had an unmodified choline headgroup (87.2 Da), for comparison with the headgroup-labeled Alexa Fluor 488 Streptavidin/Biotin-X (∼53 KDa) (FIG. 4A). Although Bodipy FL was conjugated at one of the two fatty acid tails at the C12 position of HPC, the chain length of the other lipid tail was the same as the lipid tails of other membrane components (16:0). Thus, the major difference between these membrane components resided in the soluble regions that differed in their structure and size. All three membrane components showed similar diffusion coefficients (*D*) between 0.73 and 1.1 µm^2^/s as measured by FRAP experiments (TABLE 2).

**TABLE 2.** The parameters in the two-dimensional Min dynamic model. The symbol ‘#’ indicates the number of protein molecules. *C*_*D0*_: the initial concentration of MinD in the system; *C*_*E0*_: the initial concentration of MinE in the system*; C*_*M0*_: the initial concentration of membrane component in the system.

| Parameter |  | Unit |
| --- | --- | --- |
| $C_{D0}$ | 48000 | $\#/\mu\text{m}^2$ |
| $C_{E0}$ | 55920 | $\#/\mu\text{m}^2$ |
| $D_D$ | 60 | $\mu\text{m}^2/\text{s}$ |
| $D_E$ | 60 | $\mu\text{m}^2/\text{s}$ |
| $D_d$ | 1.2 | $\mu\text{m}^2/\text{s}$ |
| $D_{de}$ | 0.4 | $\mu\text{m}^2/\text{s}$ |
| $D_e$ | 1.2 | $\mu\text{m}^2/\text{s}$ |
| $k_D$ | 0.48 | 1/s |
| $k_{de}$ | 8.65E-05 | $\mu\text{m}^2/\text{s}$ |
| $k_{dE}$ | 8.52E-07 | $\mu\text{m}^2/\text{s}$ |
| $k_e$ | 0.3 | 1/s |
| $C_{M0}$ | 5000 | $\#/\mu\text{m}^2$ |

In FIG. 4B and Movie 7, we demonstrated that the waves of both Rhodamine DHPE and BODIPY FL C12-HPC were observed in the co-localization experiments with MinE. The spatiotemporal features in the kymographs resembled those of Alexa Fluor 488-Streptavidin/Biotin-X (FIG. 4C). However, the membrane component waves were seen less frequently and no directional accumulation of Rhodamine DHPE or BODIPY FL C12-HPC was resolved from the images (FIG. 4D). Since all three membrane components had similar diffusivity, the poor ability of Rhodamine DHPE and BODIPY FL C12-HPC to form waves and establish a concentration accumulation could be attributed to the relatively small soluble regions that faced weaker steric repulsion force from the Min proteins.

### A kinetic model of the membrane component waves

FIG. 5A illustrates the proposed mechanism by which propagation of the Min protein waves can induce transport of the membrane component. When the Min protein wave propagates with time, the steric pressure at the leading edge of the protein waves increases with time, but the steric pressure towards the trailing edge of the protein waves decreases with time. Therefore, the corresponding membrane component waves are formed by the continual increase of the steric pressure behind the membrane component waves to push them forward. This is coupled with the continual release from the steric pressure in front of the membrane component waves to drag them forward. The spatiotemporal variation of the steric pressure imposed by the Min proteins causes most membrane components to locate in the region between the crests of the Min protein waves. The phenomenon resembles the linear peristaltic pumping process, by which the fluid body is delivered forward by the mechanical parts continually impose the pressure behind and release the pressure in front of a fluid body.

**FIG. 5.**
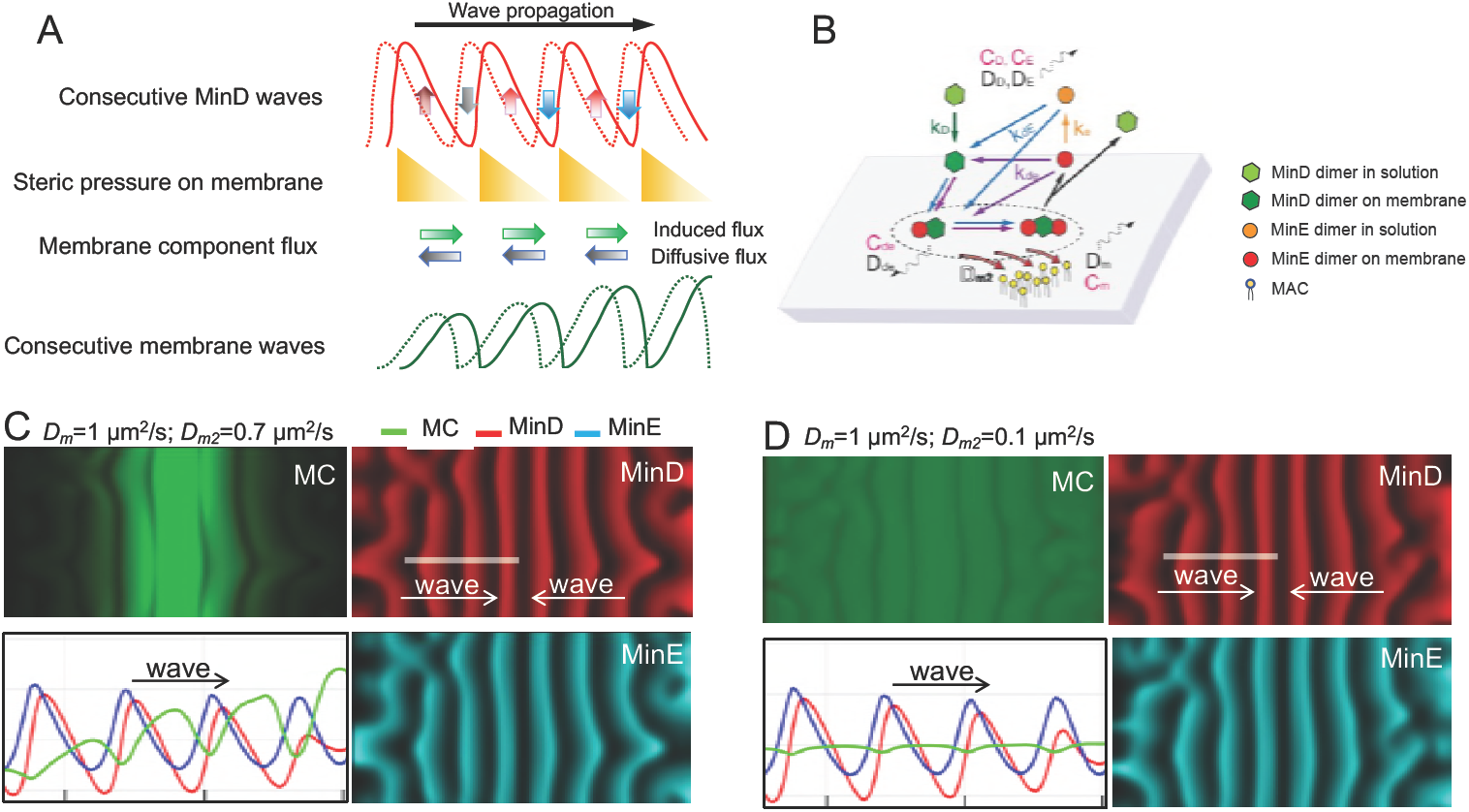
Simulated membrane component waves alongside the Min protein waves. (A) Illustration of the mechanism by which propagation of the Min protein waves induce transport of membrane components. The competition between the induced flux and the diffusive flux of the membrane component determines the concentration within a local area that further influences the overall distribution of the component in the membrane. (B) Schematic illustration explaining the interactions, rate constants, concentrations, and diffusivity factors in the numerical model. The rate constants and the corresponding steps are color coded. (C) Demonstration of how the membrane component waves can be influenced by varying *D*_*m*_ and *D*_*m2*_. The simulated waves propagate towards the center of the boxed area. The diffusivity Dm is set as 1 µm^2^/s and the repulsed ability *D*_*m2*_ is set as 0.7 µm2/s for the comparison with the experimental data using Alexa Fluor 488 Streptavidin/Biotin-X DHPE. (D) The diffusivity *D*_*m*_ is set as 1 µm^2^/s and *D*_*m2*_ is set as 0.1 µm^2^/s for the comparison with the experimental data using Bodipy FL C12-HPC. See Movie 8.

We developed a numerical model to describe the physical principles of steric repulsion that underlies the formation of the membrane component waves. An assumption that the membrane component can be expelled by MinD and MinDE complexes on the membrane was introduced to correlate the membrane component dynamics with the existing model of the Min protein dynamics (FIG. 5B; see also Materials and Methods). Therefore, lateral movement of the membrane component can be attributed not only to its own diffusion, but also to the steric repulsion caused by MinD and MinDE complexes as shown in EQ.1.

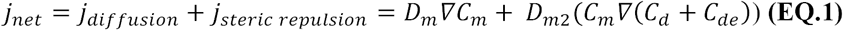

In EQ.1, *j*_*net*_ is the net flux of membrane component; *j*_*diffusion*_ is the diffusive flux caused by the concentration gradient of the membrane component; and *jsteric repulsion* is the flux induced by the steric repulsion caused by the Min protein dynamics (FIG. 5A). The diffusive flux of the membrane component due to diffusion in a fluid membrane can be described by two-dimensional Fick’s law, in which the diffusive flux is represented as the concentration gradient multiplied by the diffusivity, *D*_*m*_, of the membrane component in the fluid membrane [20, 21]. In addition, it has been shown that when the membrane surface is crowded with proteins, steric pressure can arise from collisions between proteins [22, 23]. Hence, we proposed that self-organization of MinDE within the Min protein waves can crowed the membrane plane and generate steric pressure. This steric pressure is proportional to the amount of proteins on the membrane plane, i.e. the higher the concentration of the Min proteins, the larger the steric pressure. Therefore, the concentration gradient of MinD and MinE in the protein wave can cause spatial variation of the steric pressure that provides driving force to induce the membrane component flux (FIG. 5A). In addition, the more membrane component that exists, the more that will be repulsed in a local area. Therefore, the induced flux of the membrane component is proportional to the magnitude of the driving force multiplied by the local concentration of the component. The proportionality is set as *D*_*m2*_, representing the repulsed ability of the component (EQ. 1).

### The simulated membrane component waves resembled the experimental observations

The kinetic equations were solved using the parameters listed in TABLE 2. The simulated Min protein waves showed characteristic features of the experimental Min protein waves as described earlier (FIG. 5C, D, Movie 8). Most importantly, the simulated membrane component waves corresponded with the Min protein waves and showed consistent features with the experimental observations using Streptavidin/Biotin-X DHPE (FIG. 1C, D, 3).

To explain the membrane component waves of Alexa Fluor 488 Streptavidin/Biotin-X DHPE were clearer than the Rhodamine DHPE and Bodipy FL C12-HPC waves (FIG. 4B), we proposed that the soluble regions of MinD and MinE could expel the membrane components occupying the same or vicinity space (FIG. 4A). Therefore, the membrane component with a larger soluble region could face higher steric pressure to induce a larger flux, thus associating a greater repulsed ability, *D*_*m2*_. The simulation results demonstrated that when *D*_*m2*_ was set around 0.7 µm^2^/s, the simulated component waves matched the experimental observations of Alexa Fluor 488 Streptavidin/Biotin-X (FIG. 5B). Importantly, while the Min protein waves showed consistent protein concentration, the concentration of the membrane component in waves increased following the same direction as the wave propagated, supporting the idea that the membrane component was transported along with the propagating Min protein waves. Meanwhile, when *D*_*m2*_ was decreased to 0.1 µm^2^/s to represent the repulsed ability of rhodamine and choline in the simulation, the wave amplitude and the spatial accumulation became weak, which was consistent with the experimental observations (FIG. 5C).

### The intrinsic property of the membrane component influenced its spatial distribution

Since the diffusivity (*D*_*m*_) and the repulsed ability (*D*_*m2*_) are intrinsic properties of the membrane component, we used the kinetic model to investigate different combinations of diffusivity and repulsed ability on the wave formation (FIG. 6). We studied *D*_*m*_ of 10, 5, and 1 µm^2^/s, since the diffusivity of typical lipids is in the range of 1-10 µm^2^/s [24], and the experimental diffusivity of membrane components was around 1 µm^2^/s (TABLE 3). In addition, the diffusivity of the lipid-bound peripheral membrane proteins [25] and the Min proteins (TABLE 2) [14] were reported in the same range. The *D*_*m*_ value of 0.1 µm^2^/s was also studied to mimic hypothetical components with very low diffusivity in the membrane. The repulsed ability *D*_*m2*_ was set to 1, 0.5, and 0.1 µm^2^/s, among which *D*_*m2*_ of 1 µm^2^/s was introduced to mimic hypothetical membrane components with a large soluble region that could face stronger repulsion force.

**TABLE 3.** Diffusivity of membrane components in the DOPC/DOPG (67:33 mole%) bilayer.

| Membrane component | Diffusivity ($\mu\text{m}^2/\text{s}$) | Mobile fraction |
| --- | --- | --- |
| Alexa Fluor 488 Streptavidin/<br>Biotin-X DHPE | $0.97 \pm 0.05$ (n=5) | $0.81 \pm 0.04$ (n=5) |
| Rhodamine DHPE | $0.73 \pm 0.04$ (n=5) | $0.85 \pm 0.01$ (n=5) |
| BODIPY FL C12-HPC | $1.1 \pm 0.1$ (n=5) | $1.03 \pm 0.02$ (n=5) |

<sup>a</sup> Standard error of the mean (SEM)
<sup>b</sup> The measurements were performed at different locations of each sample.

**FIG. 6.**
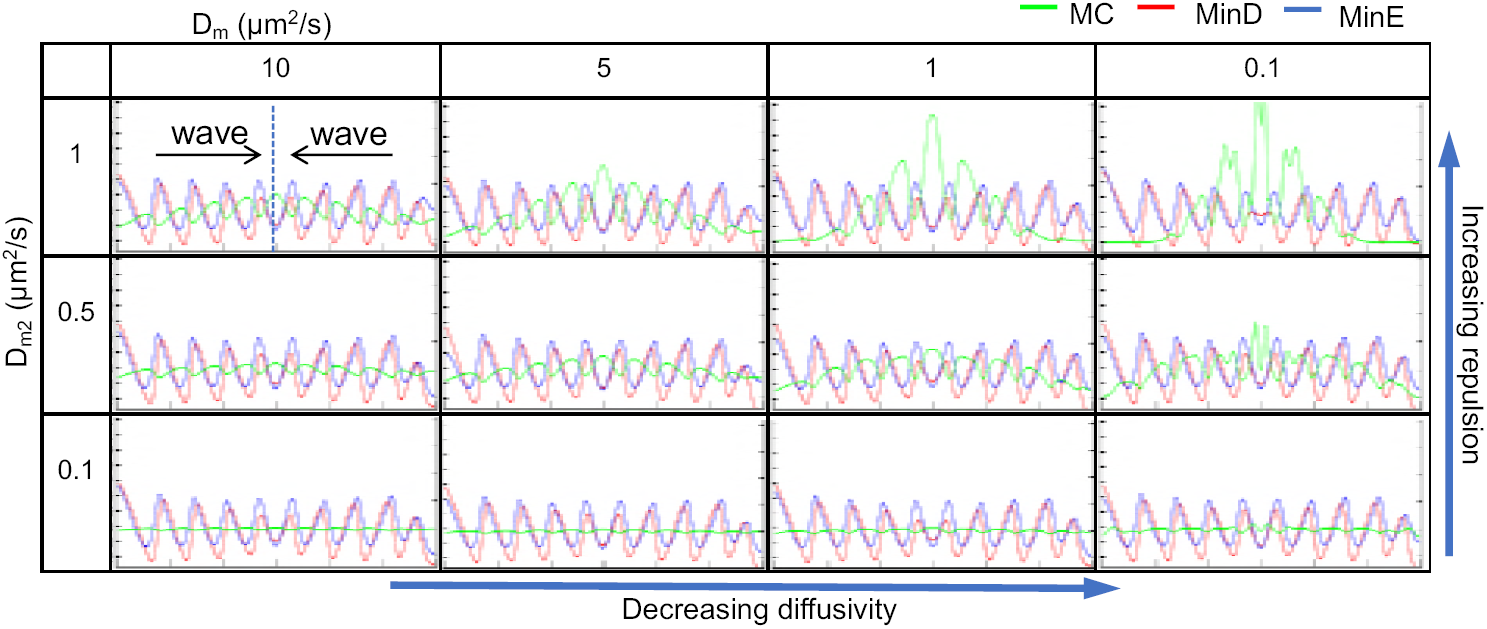
The diffusivity (*D*_*m*_) and the repulsed ability (*D*_*m2*_) of a membrane component are the critical factors contributing to the Min protein-induced steric pressure and hence membrane component waves in the membrane.

Snapshots of the simulated concentration profiles are shown in FIG. 6, which represented two opposing Min protein waves propagated from both left and right sides towards the center. Higher *D*_*m2*_ led to clearer membrane component waves when *D*_*m*_ was fixed, because higher *D*_*m2*_ caused a larger flux of the component accumulating towards the same direction as the wave propagation and led to accumulation of the component at the center where two waves collided with each other. Because the Min system exists in a dynamic equilibrium during wave propagation, the total concentration of the Min proteins in the system as well as the amount of proteins that contribute to the steric pressure gradient by molecular crowding should remain constant. Therefore, the correlation between *D*_*m2*_ and steric pressure was mainly influenced by the size of the soluble region that occupies a similar space as MinDE. While the larger soluble region faces stronger steric pressure and associates with larger *D*_*m2*_, the smaller soluble region faces weaker steric pressure and associates with smaller *D*_*m2*_. Meanwhile, since faster diffusion or increasing diffusive flux tended to counteract with the directional accumulation of the components, the amplitude of the membrane component waves and the spatial accumulation of components became weaker in corresponding with increasing *D*_*m*_ values. Taken together, because the induced flux was proportional to *D*_*m2*_ and the diffusive flux was proportional to *D*_*m*_, we thence explained why smaller *D*_*m2*_ and larger *D*_*m*_ can cause homogeneous distribution of the components, while larger *D*_*m2*_ and smaller *D*_*m*_ can cause the components to concentrate.

## DISCUSSION

To the best of our knowledge, this study reports a new mechanism that the biological reactions occurring on the membrane surface can coordinate the physical processes to transport and distribute the components in the membrane. The mechanism is demonstrated by the propagating Min protein waves formed by self-organization to actively transport the membrane components that associated with the membrane through lipid anchors. The Min protein and membrane component waves showed spatial exclusion from each other on the membrane surface. The phenomenon underlies a repulsive force between the Min proteins and the membrane component that is determined by the diffusivity (*D*_*m*_) and the repulsed ability (*D*_*m2*_) of the membrane component (FIG. 5). When the Min protein waves propagate, the time-dependent concentration change of MinDE causes the corresponding change of the steric pressure in the membrane that further induces a flux of the membrane component. When the induced flux by the steric pressure is much larger than the diffusive flux, a large amount of the membrane component can be transported along the propagating Min protein waves. This directional transport can lead to considerable accumulation of the component at the position where there is a barrier to halt transportation (FIG. 3A, 5B). The accumulation of the membrane component is also explained by the fact that MinD and MinE undergo continuous cycles of attachment to and detachment from the membrane, but the membrane component migrates passively and remains on the membrane surface during wave propagation. Therefore, the membrane component can be continuously transported by the Min protein waves and accumulate alongside the wave propagation, but the Min proteins do not accumulate on the membrane surface. The phenomenon is highlighted in the spiral waves, in which the membrane component not only showed directional accumulation, but also formed concentration gradients within the spiral wave patterns (FIG. 3). On the contrary, a strong diffusive flux of the membrane component can counteract with the directional accumulation of the component, resulting in even distribution of the component in the membrane (FIG. 6).

Since the repulsed ability of a membrane component is directly proportional to the steric property of the soluble region in the component, the Min protein dynamics may influence distribution of the component based on the size and structure of the soluble region. In another words, the stronger the steric effect of the soluble region is, the more the component can be influenced by the steric pressure induced by the Min proteins. In addition, since the diffusivity of a membrane component is mainly determined by the way that is associating with the membrane, we predict that the difference in protein structure and diffusivity of different peripheral and integral membrane proteins could tune the effectiveness of the Min proteins to induce formation of the membrane component waves. The exact correlation remains to be investigated.

In literature, counter oscillation of the Min proteins and the cytoskeletal protein FtsZ *in vivo* and anti-phase distribution of the Min proteins and FtsZ on SLBs *in vitro* were reported [26, 27], which resembled the staggered spatiotemporal distribution of the Min protein and membrane component waves reported in this study. However, direct protein-protein interaction exists between MinC and FtsZ that leads to an antagonistic effect of their membrane distribution. This mechanism is fundamentally different from the mechanism by which the membrane component waves were induced by the steric pressure imposed by the Min proteins on the membrane. Owing to the involvement of the Min proteins in the cell division process, it will be of great interest to investigate whether the Min system could transport other cell division proteins by means of steric repulsion. Finally, an intriguing implication, that arose from this *in vitro* study aiming at characterizing the transport mechanism of the membrane component by self-organization of the Min proteins, is that the Min system may function in segregating the membrane components and maintain their heterogeneous distribution in the membrane and further localize specific physiological events inside a cell.

## SUPPORTING MATERIAL

Eight movies are available.

**Movie 1**. The large-scale waterfall waves of Streptavidin/Biotin-X, MinD, and MinE. The time series was taken with a 10-second interval using the wild-field fluorescence microscope system. The frame rate is 7 fsp. This movie corresponds to FIG. 1B. Scale bar, 100 µm.

**Movie 2**. The traveling waves of Streptavidin/Biotin-X, MinD, and MinE. The time series was taken with a 6-second interval using the confocal microscope system. The video was saved at a playing speed of 7 fsp (frames per second). This movie corresponds to FIG. 1C. Scale bar, 50 µm.

**Movie 3**. The spiral waves of Streptavidin/Biotin-X, MinD, and MinE. The time series was taken with a 6-second interval using the confocal microscope system. The frame rate is 7 fsp. This movie corresponds to FIG. 1D. Scale bar, 50 µm.

**Movie 4**. The spiral waves of Streptavidin/Biotin-X in the absence of labeled proteins. The time series was taken with a 6-second interval using the confocal microscope system. The frame rate is 7 fsp. This movie corresponds to FIG. 2A. Scale bar, 100 µm.

**Movie 5**. The crawling waves of Streptavidin/Biotin-X in the absence of labeled proteins. The time series was taken with a 6-second interval using the confocal microscope system. The frame rate is 7 fsp. This movie corresponds to FIG. 2B. Scale bar, 100 µm.

**Movie 6**. Spiral waves showing membrane-component concentration gradients. The time series was taken with an interval set as freerun (estimated with a 5.26-second interval) using the confocal microscope system. The frame rate is 7 fsp. This movie corresponds to FIG. 3. Scale bar, 100 µm.

**Movie 7**. Simultaneous imaging of Rhodamine DHPE, Bodipy FL C12-HPC and MinE waves. The time series was taken with a 6-second interval using the confocal microscope system. The frame rate is 7 fsp. This movie corresponds to FIG. 4B. Scale bar, 100 µm.

**Movie 8**. Simulated wave patterns of membrane component (green), MinD (red), and MinE (cyan) on the membrane. Left panel: diffusivity *D*_*m*_ = 1 µm^2^/s, repulsed ability *D*_*m2*_ = 0.7 µm^2^/s; right panel: *D*_*m*_ = 1 µm^2^/s, *D*_*m2*_ = 0.1 µm^2^/s. The simulation time was 1000 seconds. The movie is presented with a 4-second interval and the frame rate is 33 fsp. The left and right panels correspond to FIG. 5C and 5D, respectively.

## ACKNOWLEDGEMENTS

We thank Mei-Yi Chen in the YLS lab, Meng-Ru Ho at the Biophysical core facility, and Chin-Chun Hung at the Imaging core facility of the Institute of Biological Chemistry, Academia Sinica, and Wan-Chen Huang at the Single-Molecule Biology core lab of the Institute of Cellular and Organismic Biology, Academia Sinica. This work was supported by the Ministry of Science and Technology (102-2311-B-001-020-MY3 to YLS and 105-2628-E-002-015-MY3 to LC), a Career Development grant from National Taiwan University (NTU-CDP-107L7725) to LC, and a National Taiwan University and Academia Sinica Joint Program grant (08/2015∼12/2016) and a Thematic Project grant from Academia Sinica (AS-106-TP-B04) to both YLS and LC.

## AUTHOR CONTRIBUTIONS

YLS and LC conceived, designed, performed experiments, analyzed data, and wrote the paper. LTH and BFL conceived and performed the simulation experiments. YMT and CYH conceived and performed the initial reconstitution experiments. JSC and ZXL performed the experiments. YCB, HLL, YPS, and MFH contributed to the protein production.

